# Plant-plant communication in tomato: mechanisms underpinning information transfer between plants

**DOI:** 10.64898/2026.11.03.763850

**Authors:** Claude Castella, Guy Costagliola, Doriane Dumont, Kenia Quinzoni, Anne-Violette Lavoir, Bernard Caromel, Alan Kergunteuil

**Affiliations:** INRAE, PSH, F-84000 Avignon, France; Université Côte D’Azur, INRAE, UMR ISA, 06000, Nice, France; INRAE, GAFL, CS 60094, F-84140, Montfavet, France

**Keywords:** plant-plant communication, defence priming, information theory, volatile organic compound, chemodiversity, *Solanum lycopersicum*, tomato leafminer

## Abstract

- As evidences about plant-plant communication are mounting, we lack a mechanistic understanding of intricate events occurring inside and among pairs of plants. This is needed to untangle proximal causes explaining how volatiles emitted by herbivore-attacked plants can enhance plant defence in intact neighbouring plant. A comprehensive knowledge of these interactions may favour their integration in trait-based agroecological frameworks.
- Based on an original experiment set-up, we connected infested sender plants and intact receiver plants during 3 days. We monitored metabolomic reprogramming of sender plant and progressive adjustments in their volatile emissions. Plant-plant communication was tested according to metabolomic priming of receiver plant. Information transfer was tracked with kinetics of volatile profiles and chemodiversity metrics grounded in information theory.
- Our results show that sender plants infested with Phtorimae absoluta undergo deep reconfiguration of immune systems and balance their emissions of mono- and sesqui-terpenoids over 3 days. These changes in airborne cues result in similar reprograming of immune system in receiver plants. Additionally, we observed herbivory-driven relationships between chemodiversity of volatile profiles produced by sender and receiver plants.
- Both metabolome and volatilome suggest that plant-plant communication operates as mirror effects with receiver plants matching up their defences on that of sender plants. From an adaptative standpoint, this supports the idea that information conveyed among plants is explicit enough to warn receiver plant against appropriate stress.
- More generally, this study points that information transfer could rely not only on composition but also on chemodiversity of volatile cues. We discuss implications related to cue detection at the molecular scale and cue propagation at the plant cover scale. Bridging composition and chemodiversity of volatile cues opens interesting avenues to study the efficiency of communication system in noisy environments.

## 1. Introduction

Plants process multiple cues from their environments (Karban, 2021). Among them, volatile organic compounds (VOC) draw attention, and sometimes imagination, as airborne cues mediating plant-plant communication (Schuman, 2023). In plant ecology, experiments conducted in natural conditions are fundamental to gauge their prevalence (Hagiwara et al., 2024; Pearse et al., 2013). They also bring insights in biodiversity-ecosystem functioning relationships (Medina-van Berkum et al., 2026). In parallel, four recent reviews highlight the need to address gaps in knowledge about mechanisms supporting plant-plant communication (Brosset and Blande, 2022; Kessler et al., 2023; Ninkovic et al., 2021; Schuman, 2023). This is of valuable interest to move from empirical and experimental evidences to functional frameworks addressing a series of events leading to plant-plant communication.

Research about plant-plant communication has long revolved around herbivore-induced plant volatiles (HIPV) emitted by plants in response to insect attacks (Guerrieri, 2016). Since these compounds can elicit defences in intact neighbouring plants, they offer interesting avenues to favour pest control at the crop scale from the earliest stages of pest infestation. In this context, we take advantage of tomato as a highly chemically-diversified model to investigate mechanisms of plant-plant communication (see Amegan et al., 2024 for volatile diversity; Exonam Amegan et al., 2026 for leaf metabolism diversity). Despite clues suggesting that domestication has affected the diversity of volatile profiles (Amegan et al., 2024), a body of literature indicates that modern cultivars of tomato can still induce sesquiterpenoid emissions when attacked by a major pest like the tomato leafminer, Phtorimae absoluta (e.g., Ayelo et al., 2022; Chen et al., 2026). Interestingly, sesquiterpenoids have been hypothesized to convey large information bandwidth according to their high structural diversity (Rosenkranz et al., 2021). Parallel to herbivore induction of sesquiterpenoids, four studies have evidenced plant-plant communication in tomato based on synthetic volatiles (Pérez-Hedo et al., 2021) or sender plants infested with Macrosiphum euphorbiae (Catola et al., 2018) or Spodoptera spp. (Coppola et al., 2017; Sugimoto et al., 2014). Following these proof-of-concepts about plant-plant communication in tomato, we think that chemical ecology offers a sound mechanistic approach to integrate a three-step process involved sequentially in (i) cue production by sender plants, (ii) information transfer between plants and (iii) response of receiver plants. For this purpose, we combine chemical ecology with metabolomic and information theory.

First, recent analytical breakthroughs are particularly promising to study metabolomic reprogramming related to plant immune systems (Williams et al., 2021). On the specific side of sender plant, bringing together untargeted metabolomic of plant tissues and volatile analyses could help to explore proximal causes explaining the production of airborne cues. On the side of receiver plant, metabolomic is increasingly used to test responses to these volatile cues (e.g., Ameye et al., 2020; Ceccanti et al., 2025). Furthermore, if we consider metabolism as a mechanistically-explicit repository of traits that are fruitful to predict plant functioning (Walker et al., 2022), we can expect that untargeted metabolomic of receiver plants may help to determine the information they process before to adjust their response.

Second, most of the attention paid to informative transfer among plants have been restricted to descriptive approaches reporting compositional changes of volatile profiles. In this context, bridging chemical ecology and information theory could help to better understand how information is encoded. Pioneer works have suggested that the presence of particular compounds is critical in relatively poor signals (de Moraes et al., 1998; Doyle, 2009). However, as analytical developments have expanded the inventory of potential infochemicals, mathematical descriptors grounded in information theory became pivotal to link chemodiversity with information amount (Li et al., 2020). In this regard, recent work has been done to develop appropriate metrics of chemodiversity integrating metabolomic richness, evenness and structural disparity (Petrén et al., 2024, 2023). These new metrics of chemodiversity are promising to decipher informative channels supporting plant-plant communication. This line of research is important given that, so far, most of studies interested in the interplay between information amount and the richness of cues have addressed plant interactions with fungus, herbivore or parasitoid (Hilker, 2014; Petrén et al., 2024; Raguso et al., 2015).

Here, we investigate plant-plant communication based on an original two-step experiment accounting for a “memorisation” delay during which receiver plants are no longer exposed to informative cues. Thereby, we aim at testing priming processes sensus stricto (Martinez-medina et al., 2016). More particularly, we address three questions related to three intricate events leading to plant-plant communication. First, we explore how insect attacks on tomato reconfigure plant metabolome and emissions of HIPVs as source of cues. Second, we test if cues perception in receiver plants result in metabolomic reprogramming. Finally, we investigate how chemodiversity of VOC profiles released by sender plants helps to understand information transfer among plants.

## 2. Material and Methods

### 2.1. Plant material

Seeds of the tomato accession ‘Rose de Berne’ were obtained from the biological resource centre “CRB-leg” hosted by INRAE (Salinier et al., 2022) and plant growing was planned in greenhouses managed by the experimental unit ‘A2M - Unité Expérimentale Arboriculture et Maraîchage Méditerranéens’ (INRAE France, Avignon). Plants were sown in small pots (7×7 cm) filled with seedling substrate (Klassman; N:P:K 12:14:24 kg.m-3). After one week, seedlings were potted in larger pots (13×13 cm) with clay substrate (Klassman; N:P:K 14:10:18 kg.m-3). The preparation of plant material was repeated weekly between May and June 2025. This way, we were able to conduct the experiment between June and July 2025 with 4-weeks-old plants at a vegetative stage with analogous biomasses.

### 2.2. Insect material

Insect rearing was done at the insectarium of the research unit ‘PSH-Plantes et Systèmes de Culture Horticoles’ (INRAE France, Avignon) in a room with controlled conditions (20±0.5 °C, 60%±10 % RH). Natural light was supplemented by neon lighting set with a photoperiod of L16:D8. In April 2025, we initiated a rearing of Phtorimae absoluta with larvae collected in south of France (Vaucluse, Bouche-du-Rhône). For rearing purpose, the reproduction of tomato leafminer was done with a set of 2 tomato varieties (i.e., Momor, Marmande) purchased from Vilmorin (Paris, France). For experimental purpose, larvae were retrieved weekly from age-controlled cohorts. In brief, plants were exposed to gravid females for 48h before isolation from any insects until egg hatching (i.e. 6 days after oviposition). Young larvae with 2-days old maximum were teased apart with a scalpel under binocular. Thereby, we were able to control plant infestation with the first larval instar.

### 2.3. Experimantal set-up

Experiments were conducted in a room with controlled photoperiod (L16:D8), temperature (L:D 22°C:18°C) and relative humidity (60%). Sender and receiver plants were enclosed in individual gall jars (20 x 50 cm; c.a. 15 L) connected by PTFE tubing (Fig. 1). Airflow was drawn into the system from the sender to the receiver plants using a diaphragm pump (KNF, Laboport, 10 L.min-1). At the entrance of the system, cleaned air was obtained with a charcoal filter. The experimental set-up relies on two types of pairs of plants. In one case, intact sender plant were associated to intact receiver plants (i.e. control with constitutive VOC as airborne cues). In the other case, sender plants were infested with 6 larvae (L1) of P. abosulta while receiver plants remained intact (i.e. stress treatment with HIPV as airborne cues). Three randomized pairs of plants were tested weekly on 6 consecutive weeks. In total, 9 pairs of plants for each modality ‘control’ and ‘stress treatment’ were tested. Experimental design was based on two-step time-series (Fig. 1).

**Figure 1:**
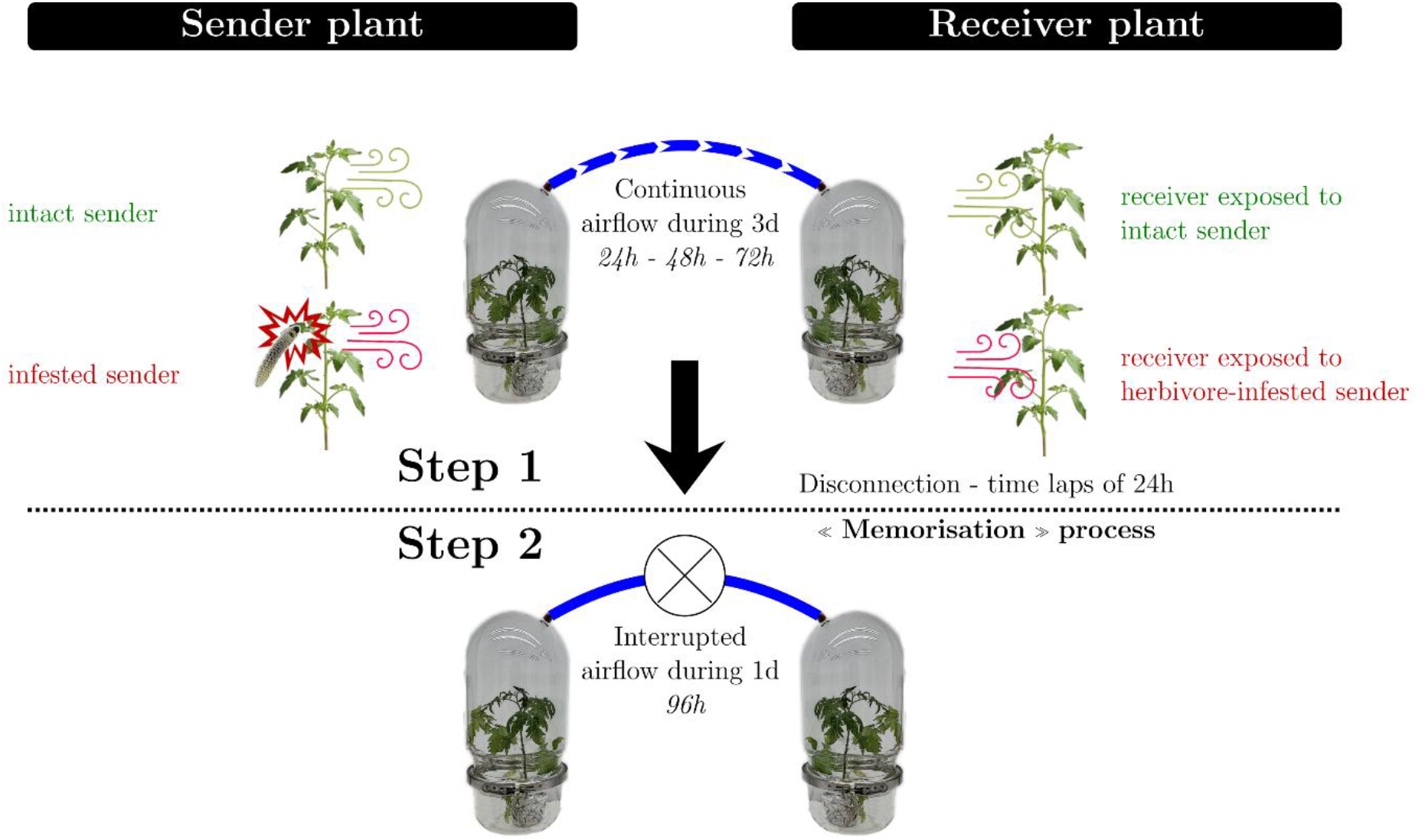
Two-step experimental setup. During step 1, air was transferred from glass jar with sender plant to glass jar with receiver plant. Step 2 started 3 days after with the interruption of air transfer to isolate receiver plant from airborne cues produced by sender plant. Priming of receiver plant was tested 4 days after the beginning of the experiment.

In the first step, sender and receiver plants were connected each other to allow for information transfer. Volatile released by each plants were collected at 24h, 48h and 72h. In the glass jars with receiver plant, a set of taps was used to purge during 20 min the VOC released by sender plants before each volatile collection. In the second step, after 72h, sender and receiver plants were disconnected. Sender plants were sampled for metabolomic and the receiver remained supply to a pure air flow for a further 24 hours. After this period, we conducted final VOC collection and metabolomic sampling of receiver plants.

### 2.4. VOC collection and GC-MS analyses

Plant volatiles were collected on TenaxTM TA cartridges (200 mg of adsorbent phase, 60/80 mesh size) that were pre-conditionned at 200°C for 1h on a sorbent tube conditioner (TC-20 Markes international) with a nitrogen airflow (50 mL.min-1). Volatile collection was performed during 20 min with a pump settled at 200 mL.min-1. After VOC collection, cartridges were stored at 4°C for two weeks maximum before analytics.

Before injection, cartridges were spiked with 40 ng of internal standard (azulene, Sigma-Aldrich) with an adsorbent tube injector system (ATIS, Supelco, 1µL injected at 100°C for 2 min, airflow at 30 mL.min-1). The cartridges were analysed on a Trace GC-MS ultra/ISQ system (single quadrupole; Thermo Scientific) combined with a Turbomatrix–650ATD thermal desorber (Perkin Elmer), using an DB5-MS column (30 m × 0.25 mm × 0.50 μm; Perkin Elmer). The first VOC desorption from the cartridges to the cold trap was achieved by heating the cartridges to 280 °C for 10 min. The second desorption from the cold trap to the GC was achieved with a flash-heat at 280°C for 5 min. In the GC, the temperature program was set as following: 2 min at 40 °C, first heating ramp to 100 °C at a rate of 20 °C per min, second heating ramp to 190 °C at 5 °C per min, and finally to 300 °C at 30 °C per min. Mass detection was done with electron impact (70eV) for ions ranging from 43 to 500 m/z.

Chromatogram deconvolution was done with the R package ‘`xcms’` (Smith et al., 2006), associated with ‘`MsExperiment’` (Gatto et al., 2024) and ‘`MsFeatures’` (Rainer, 2024) available from the Bioconductor community (Morgan and Ramos, 2025). The R script used for deconvolution is available in Supplementary Material and the chromatograms have been uploaded in the public data portal ‘Recherche Data Gouv’ (https://doi.org/10.57745/V5OCJ2). Peak detection was conducted from a list of 24 ions with distinct m/z values specific to alkane and green-leaf volatiles (43, 57, 71, 77, 79, 85, 87 m/z), terpenoids (68, 69, 91, 92, 93, 119, 120, 121, 134, 136, 204 m/z) and phenylpropanoids (104, 105, 107, 128, 152, 161 m/z) that were eluted between 5 and 24 min of retention time. In total, 30 687 peaks were detected with the ‘centWave’ algorithm after we optimized parameters as following: peak width between 1 and 10s, resolution at 250 ppm, signal/noise threshold at 10 and mzdiff at -0.5. Peak alignment was performed from a sample with intermediary position in the injection list and harbouring relatively high concentration of monoterpenes, i.e. an infested sender plant (mean correction time=1.41 s). Correspondence analysis based on peak grouping across samples (minimum fraction of 50% within sample groups) resulted in 143 features that were subsequentially integrated with the recommended ‘ChromPeakAreaParam’ algorithm for gap filling. Further, the compounding approach developed by (Rainer, 2026) was used to group ions belonging to a same molecule into a pseudo-spectra. In doing so, we grouped features into retention time window (10s) and, then, we teased apart co-eluted features based on Pearson’s pairwise correlation (threshold at 0.7). As a result, we obtained 126 pseudo-spectra for which only base peaks were retained for further analyses. Each of them was manually curated to remove contaminants originating from the analytical system (e.g. siloxane derivates), those found in ambient air (i.e. volatiles detected in empty glass jars) and those specific to n-alkane standards. Finally, we obtained a dataset with 14 VOCs that were putatively annotated with comparisons of mass spectra (NIST database 3.0) and experimental Kovat’s indices (C7-C20 n-alkane). Six VOCs were strictly identified with the injection of the respective standards purchased from Sigma-Aldrich (Saint Louis, USA): 2-buty-hexanol, α-phellandrene, α-terpinene, methyl salicylate, β-caryophyllene, humulene.

### 2.5. Metabolomic sampling and UHPLC-ESI-MS/MS analyses

Plant tissues were collected on the two terminal leaflets from the two youngest leaves fully developed. Samples were flash-frozen in liquid nitrogen and stored at -80°C until grinding. The whole procedure of sample preparation before injection was conducted at cold temperature to favour biochemical integrity of metabolome. In this line, grinding procedure in liquid nitrogen with a ball mill was conducted as closely as possible to extraction. Sample solubilization was done in microtubes maintained in cold supports (-20°C) with 50.78 ±0.92 mg of plant powder supplemented by 500µL of cold mix MeOH:H_2_O (80:20). Then, cell wall was broken during 1 min with cold ultrasonic bath before sample mixing on a laboratory rotator (20 min) at 4°C. Finally, samples were centrifugates (10 min at 4°C and 13,200 rpm) and filtered according to size exclusion (PTFE filters, 0.2 µm). Lastly, 199 µL of supernatant were transferred in injection vials and supplemented with 1µL of internal standard (taxifolin, 25 µM final concentration) before injection into the UPLC-ESI-QTOF systems (Waters Acquity class I, Synapt G2-Si).

Separation was carried out in an ACQUITY UPLC BEH C18 column (50 × 2.1 mm, 1.7 μm particle size) with a BEH C18 pre-column (5*2.1 mm, 1.7 µm particle size) (Waters). The column was kept at 45 °C whereas the autosampler was set at 10 °C. The injection volume was 1 μL. The sampling list was divided in batches that were randomized in their injection order. Each injection batch contained a system blank to correct for instrumental contamination, an extraction blank to clean the sample and a QC sample (mix of each extract). The initial gradient was 99 % mobile phase A (water 0.1 % formic acid) and 1% mobile phase B (acetonitrile 0.1 % formic acid). The flow rate was 0.4 mL/min. For 1.5 minutes, the gradient remained stable; then, over 15 minutes, the gradient gradually shifted to 1% of the mobile phase A and 99% of the mobile phase B. This plateau was maintained for 2.25 minutes before returning to 99 % of mobile phase A for 1 minute to allow the column to re-equilibrate prior to the next injection. The ESI source was used in negative ionization modes. Data acquisition was performed DDA mode with 5 most ion in resolution mode and the use of an ion exclusion list. Leucine Enkephalin (1 ng/µL in 50% acetonitrile 0.1% formic acid) was used as a lock spray mass reference for continuous mass correction during QTOF acquisition. The settings to the negative ion mode were: capillary voltage 1 kV, source temperature 100 °C, desolvation temperature 400 °C, cone gas-flow 50 L/Hr, desolvation gas-flow 800 L/h, sampling cone 40 V and trap collision energy 4 eV. Scan mass range was 50 Da to 1200 Da to MS mode and 50 Da to 1000 Da to MS/MS mode. Masslynx 4.2 chromatographic software (Waters, Manchester, UK) was used for data visualisation in parallel to data acquisition.

Chromatograms obtained in LC-MS were deconvoluted with the same R packages used for GC-MS. Additionally, dereplication was done with three supplementary packages developed by Rainer et al., (2022) – ‘`Spectra’`, ‘`MetaboAnnotation’` and ‘`MetaboCoreUtils’`, in combination with ‘`AnnotationHub’` (Morgan and Sheperd, 2024). The R script used for deconvolution and dereplication is available in Supplementary Material and the chromatograms have been uploaded in the public data portal ‘Recherche Data Gouv’ (https://doi.org/10.57745/V5OCJ2). Contrary to GC-MS, an untargeted approach was used for peak detection after we optimized parameters of the ‘`CentWave’` algorithm as following: peak width between 2 and 10 s, resolution at 10 pm, signal/noise threshold at 5. To avoid artefact detection, features were pre-filtered to keep only those with 3 consecutive peaks above 1e4 intensity. For LC-MS, because the ‘`CentWave’` algorithm can generate multiple peaks for broad peaks or peaks with irregular shapes, we refined the 36818 peaks detected as recommended by (Louail and Rainer, 2026). Briefly, we merged peaks that overlapped in a 4 s retention window per file and for which the signal between them is lower than 75% of the smaller peak’s maximal intensity. As a result, we obtained 23577 refined chromatographic peaks. They were aligned relatively to a pool of samples used as a quality control injected at an intermediary position in the injection list (mean correction time of 4.93 s). One sample displaying irregular alignment (accumulated correction time of 152 s) was removed from further analyses. Peak grouping was done with similar parameters to GC-MS and resulted in 651 features. No gap filling was conducted on LC-MS to limit the risk of misintegration. In general, gap filling compensates missing peak detection from noisy chromatograms or low intensity signals (Louail and Rainer, 2026). The compounding approach used to group ions belonging to single molecules into pseudo-spectra (see above for GC-MS; Rainer, 2026) was performed on retention time windows of 5s and Pearson’s correlation threshold of 0.7. Further analyses were conducted with the 499 base peaks retrieved from the resulting pseudo-spectra. These features were curated automatically by removing those with a ratio signal/blank < 30. Consequently, only 295 features were considered as plant signals. The putative annotation of these features was performed with MS² and, then, MS^1^. First, 22 features were associated with 88 spectra of MS². We retained the 22 spectra with the highest TIC intensity for spectra comparison with the MassBank database available with the Bioconductor AnnotationHub web resource (release 2024.11; 122,512 MS² spectra). In brief, experimental and database spectra were cleaned by removing low-intensity peaks (<5% of major peak) and intensity was normalized between 0 and 100 before pairwise comparison spectra based on the weighted entropy score developed by (Li et al., 2021) and following recommendations of (RforMassSpectrometry et al., 2026) with pre-filtering to account for precursor ions and 30 ppm of tolerance for m/z matching of fragmented ions. Seven features matched 91 spectra from database. The assignation of FT326 to dronedraone (i.e., a synthetic drug) was considered as an uncohesive annotation. For each of the six remianing features, we retained the annotation with the highest entropy score (ranging from 0.73 to 0.95; Tab. S1; see Fig. S1 for pairs of spectra as mirror plots). For the other features, putative annotations were performed with comparisons of experimental masses and 792,364 exact masses available from the taxonomically-explicit Lotus database (Rutz et al., 2022, 2023). Mass comparisons were conducted based on 10 ppm of maximal mass-dependant differences with a single adduct, i.e. [M-H]^-^, to limit the risk of uncohesive annotations. In total, 200 metabolomic features matched 23,454 exact masses. For each feature, hits associated with compounds already found in the *Solanaceae* family were priorly kept before filtering based on ppm error. Consequently, 23 % of the annotations concerned compounds previously described in the *Solanaceae* and ppm error ranged from 0.0005 to 9.7783 (Tab. S1). Finally, each feature assigned to an InChIKey and a SMILES, whether from MS² or MS^1^ annotations, was categorized with the R package ‘`chemodiv’` (Petrén et al., 2023) into chemical classes according to the *NPClassifier* ontology (Kim et al., 2021).

### 2.6. Statistical analysis

Statistical analysis were carried out using `R` `4.4.1` (R Core Team, 2024) with the nine following packages : `chemodiv` (Petrén et al., 2023), `igraph` (Csárdi et al., 2026), `vegan` (Oksanen et al., 2025), `caret` (Kuhn and Max, 2008), `car` (Fox and Weisberg, 2019), `lme4` (Bates et al., 2015), `merTools` (Knowles and Frederick, 2026), `rcompanion` (Mangiafico, 2026) and `multcompView` (Graves et al., 2024). The full reproducible code is available in Supplementary Materials.

The intensities of metabolomic feature detected in LC were standardized by the fresh biomass used for extraction and an internal standard displaying a good detection rate in negative mode (see 2.5 : taxifolin, C_15_H_12_O_7_, m/z=303.0499 as [M-H]^-^). Accordingly, metabolic features were given as ng (taxifolin equivalent).g^-1^. Four features displaying near-zero variance (proportion of distinct values > 90%) were removed from further analyses. In total, 291 features were used as input for data analysis. In the same line, VOC emissions were standardized by fresh plant biomass, capture duration and internal standard (azulene, C_10_H_8_, m/z=128 as base peak) and were given in pg.(azulene equivalent).g^-1^.h^-1^. Prior data analysis, the concentration of metabolic feature and VOC emissions were pre-treated with pareto scaling as a best compromise between the normalization of peak areas, the equalization of feature weighting and the integrity of original data (Van Den Berg et al., 2006).

As a first step of metabolomic data analyses, we used molecular network to map out the breadth of chemodiversity we detected. For this purpose, we computed compound dissimilarity based on PubChemFingerprint. This structure-based encoding system retrieved from SMILES and InchiKey accounts for 881 Boolean bits categorized in 7 sections such as element counts, rings or atom pairs (Boldini et al., 2024; National Center for Biotechnology Information, 2009). Edges of molecular network were determined with similarity score>0.75. Significant differences between metabolome of sender plants (i.e., intact *versus* infested plants) and receiver plants (i.e., plants exposed to intact VOC *versus* HIPV) were tested at two levels. First, for each chemical class, we conducted a Wilcoxon rank sum test based on the sum of relative intensities of metabolites. Second, for each individual metabolites, we plotted log_2_ fold-change relatively to significance of Wilcoxon test (aka volcano plot) to identify compounds contributing to metabolomic reprogramming.

To track for informative cues within VOC profiles of infested sender plant we first used a graphical approach. On a heatmap, we displayed compound-specific changes of VOC between intact plants and herbivore-infested plants (log_2_ fold-change). Statistically, we used one-tailed Wilcoxon rank sum test to estimate significant differences at 24, 48h and 72h. Further, for receiver plants, we tested if exposure to intact or herbivore-induced volatiles affected VOC emissions at 24h, 48h, 72h and 96h according to a combination of unsupervised and supervised analyses. While the unsupervised analysis was thought as *a posteriori* exploratory approach centred on patterns of VOC composition, the supervised analysis provided *a priori* predictive approach based on computation integrating larger properties of VOC profiles, including not only multi-variate composition but also hidden partens like richness, evenness or disparity. The unsupervised analysis consisted in a non-metric multidimensional scaling. Briefly, we run 5 NMDS with different computation method of dissimilarity indices to select the one giving the lowest stress value, i.e., the Euclidean index. We completed the NMDS ordination with statistical tests according to permutational multivariate analysis of variance (aka permanova) using Euclidean distance matrices. For the supervised approach, we conducted a linear-discriminant analysis based on VOC matrix of receiver plants as inputs to predict if these plants were exposed to intact volatiles or HIPV. For this purpose, we split total datasets into train and test datasets according to a stratified partitioning, 80:20 respectively. This way, we were able to estimate correct classification rates of LDA as a proxy of predictive power of VOC profiles. Due to stratified partitioning, 100 bootstrapped simulations were done to cover the solution space of possible predictions. This enabled to calculate uncertainty of correct classification rates.

Finally, we explore links between the chemodiversity of VOC profiles released by sender plants (i.e. source of cues) and the chemodiversity of receiver plant, focusing on metabolomic or VOC profiles whether we were interested in the output of plant-plant communication (i.e., metabolomic fingerprints of receiver plant collected at the end of the experiment) or signal propagation (i.e., kinetic of VOC profiles released by receiver plants all along the experiment). To do this, we first estimated for each metabolomic and VOC profiles compound dissimilarity matrices based on pairwise Jaccard distances between PubChem fingerprints of individual compounds. Second, we computed chemodiversity as abundance-weighted functional Hill diversity index with diversity order q=1 as recommended by Petrén et al. (2024). We used modelling approaches to test the relationships of chemodiversity within specific pairs of sender and receiver plants. Linear models were used to test the links between volatile chemodiversity of sender plants and metabolomic chemodiversity of receiver plant sampled at the end of the experiment (i.e., 3 days of exposition to sender plants followed by 24h of memorisation). To test the links between volatile chemodiversity of sender and receiver plants all along the experiment, we used mixed linear models. Thereby, we accounted for time series of VOC collection and we implemented in the model pairs of plants as as random effects.

## 3. Results

### 3.1 How insect attacks reconfigure airborne cues produced by sender plant ?

#### 3.1.1 Metabolomic reprogramming of leaf content relies on amino acids and terpenoids

Although 33% of metabolites were not classified (Tab. S1), our results indicate that metabolomic composition of intact sender plant was dominated by terpenoids (22%) followed by amino acids and peptides (11%), shikimates and phenylpropanoids (10%) and alkaloids (8%).

These proportions about qualitative composition mirrors relatively well the relative quantities of the different chemical classes (Fig. S1).

As compared to this basal metabolome, we observed a metabolomic reprograming of infested sender plant (Fig. 2). Overall, as show in Fig. S2, infestation resulted in 37% increase of terpenoids (one-tailed Wilcoxon test, W=16, p.value=0.052) and 20% increase of amino acids and peptides (one-tailed Wilcoxon test, W=15, p.value=0.041). At the compound level, 18 metabolites differed significantly between intact and infested sender plants (Fig. S3). Among them, we found 4 terpenoids (FT489, FT238, FT530, FT276), including an apocarotenoid (FT238) which was previously described in *S. lycopersicum*. In addition, herbivore infestation affected the synthesis of 4 amino acids and peptides (FT294, FT555, FT611, FT394), 3 fatty acids (FT169, FT196, FT332) and 1 alkaloid (FT118) (Fig. S3).

**Figure 2.**
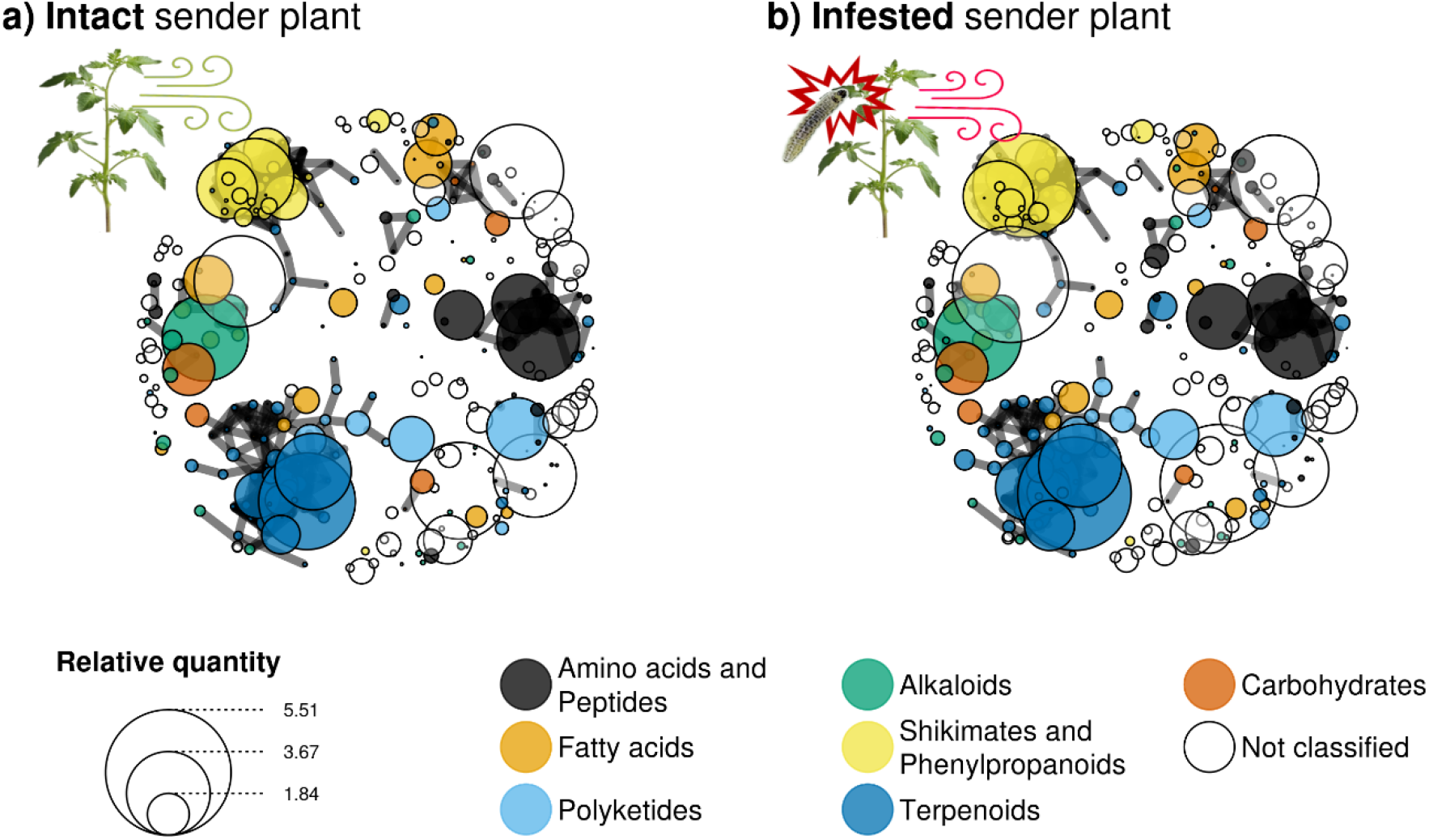
Molecular network of sender plants. The 291 vertices display the metabolomic features detected. The 651 edges between vertices were computed based on PubChem fingerprint (similarity score >0.75).

#### 3.1.2 Progressive adjustment of herbivore-induced volatiles results in a dynamic of informative cues

Constitutive emissions of intact sender plant were mainly composed of 4 monoterpenoids: α-phellandrene, 2-carene, α-terpinene and β-phellandrene (Fig. 3). Only traces of sesquiterpenoids were emitted. In infested sender plants, volatile profiles are even more dominated by monoterpenoids after 24h of herbivory. Three monoterpenoids exhibited a clear increase in emission rates: 51% for 2-carene, 42% for β-phellandrene and 16% for α-terpinene. After 48h of herbivory, emissions rates were more erratic. However, 72h after infestation, herbivory resulted in an opposite pattern with an over-expression of sesquiterpenoids (65% of increase) and a down-regulation of monoterpenoids (70% of reduction) as compared to intact sender plant. The sesquiterpenoid with the sharpest increase is β-caryophyllene. After 72h, infested plants emit 10 times more of this compound as compared to intact plants.

**Figure 3:**
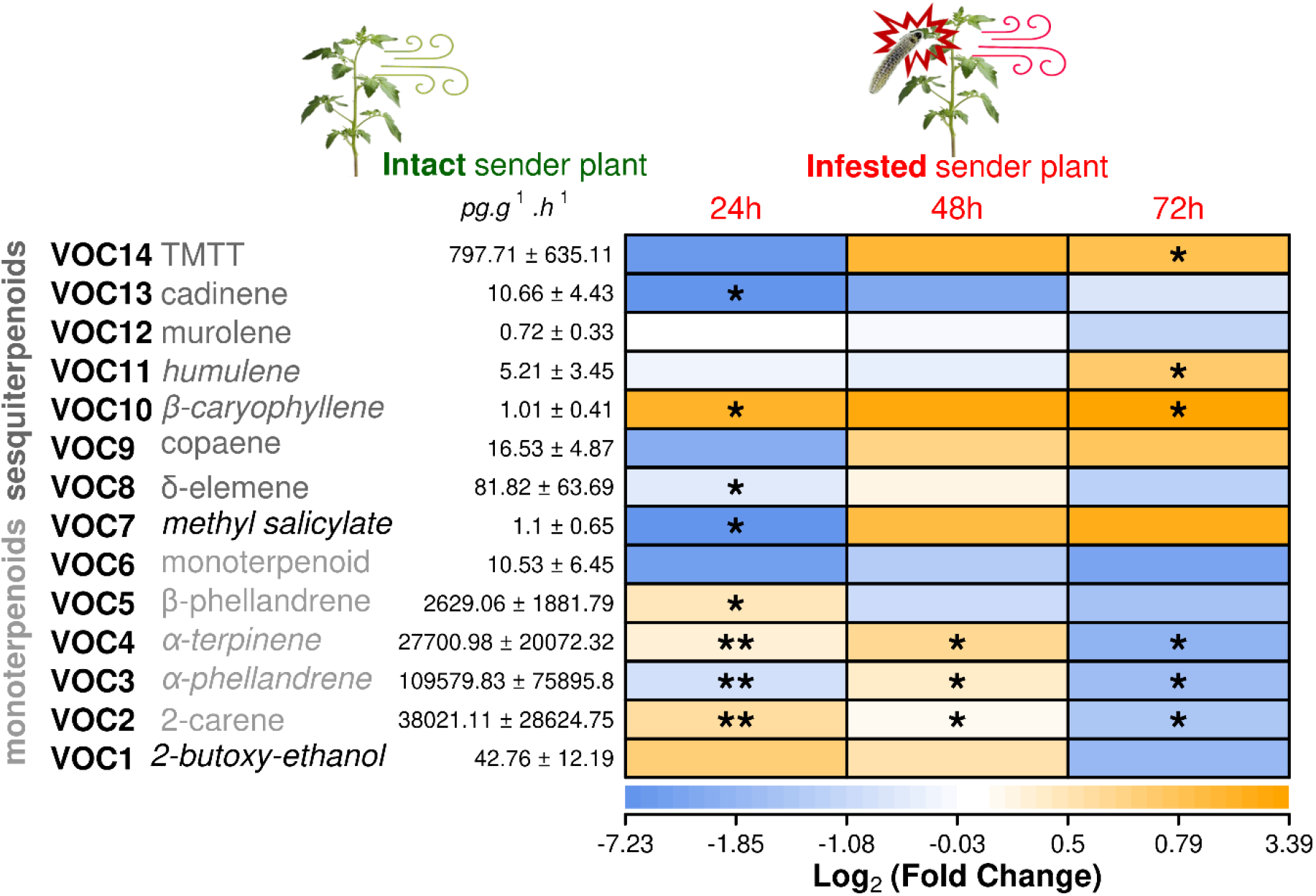
Quantitative fold change of volatile cues emitted by sender plants. Compounds in italic were identified with standards. Cold and warm colors display down- and up-regulation of compounds released by infested sender plant as compared to intact, respectively. Asterisk indicates significant differences in emission rates of individual volatile between intact and sender plant (one-tailed Wilcoxon rank sum test; * : p.value <0.05; ** : p.value <0.01)

### 3.2 Do receiver plants perceive signals released by herbivore-infested sender plants?

#### 3.2.1 The metabolome of receiver plant is reprogrammed

Receiver plants exposed to HIPV emitted by herbivore-infested sender plants down-regulated 6 features (FT064, FT481, FT347, FT345, FT017, FT262) as compared to receiver plant exposed to intact sender plant (Fig. 4). In opposition, exposure to HIPV resulted in the up-regulation of 28 metabolites (Fig. 4). In addition to 4 amino acids and peptides (FT605, FT293, FT555, FT571), they include 2 fatty acids already detected in *Solanaceae* (the dicarboxylic acid FT026 and the glycerophospholidi FT265) and 1 conjugate of jasmonic acid (FT196). Furthermore, we reported a significant increase of specialized metabolites such as shikimates and phenylpropanoids (the quercetin-3-O-rutoside FT354 and a glycosylated flavonol FT253 previously found in *Solanaceae*). Although 2 alkaloids were also up-regulated (FT172, FT560), terpenoid is the chemical class that was the most affected by previous exposure to HIPV. Indeed, 8 terepenoids shown an increase: 3 sesterpenoids (FT252, FT270, FT276), 2 triterpenoids (FT240, FT295) and 3 tetraterpenoids (FT373, FT530, FT568).

**Figure 4:**
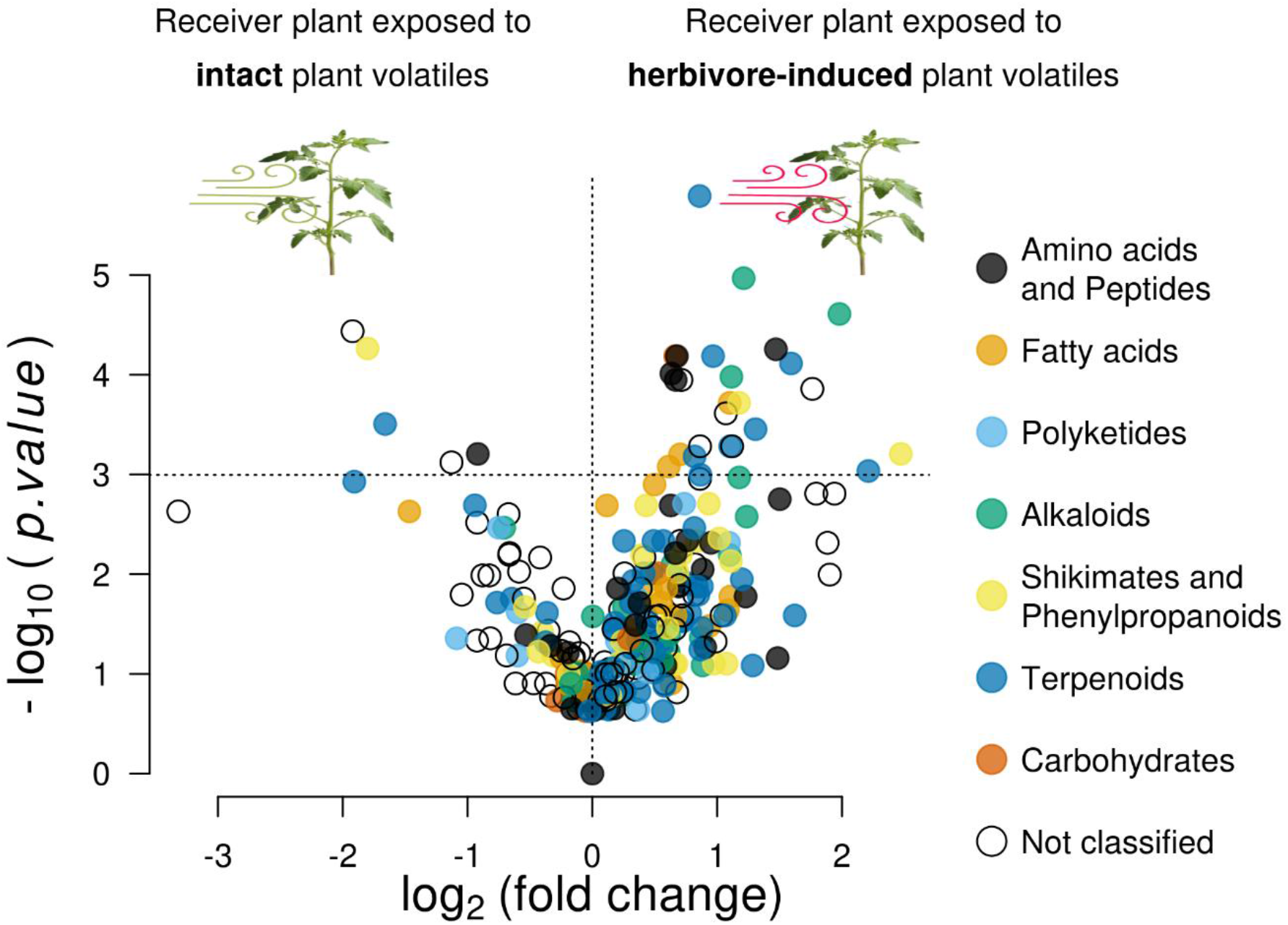
Metabolomic reprogramming of receiver plants. The 291 metabolomic features detected are coloured according to chemical pathway (NPCclassifier ontology). The vertical line separates feature over-expressed in receiver plant exposed to herbivore-infested sender plants (positive values on x-axes) as compared to intact sender plants (negative values on x-axes). Features with significant differences are displayed above the horizontal line (one-tailed Wilcoxon rank sum test; p.value < 0.05).

#### 3.2.2 The volatile profiles of receiver plants help to predict exposition to intact or herbivore-infested sender plants

The composition of volatile profiles released by receiver plants was relatively similar whether they were exposed to intact or herbivore-induced sender plants. Graphically, the unsupervised approach failed to point compositional differences of VOC profiles at 24h, 48h, 72h or 96h (Fig. 5a). In addition, we conducted a supervised approach based on a linear-discriminant analysis (LDA) to test if VOC profiles emitted by receiver plants can predict previous exposure to intact or herbivore-infested sender plants (Fig. 5b). While the unsupervised analysis is based on multivariate composition of volatile profiles considered as a mere juxtaposition of independent VOC, the supervised analysis may help to capture latent group-level properties of volatile profiles related to chemodiversity, like richness, evenness or structural disparity. At 24h, prediction accuracy of LDA was similar to random classification (49±4%). However, as long as exposure of receiver plant to intact or herbivore-infested plants increased, we recorded a gradual improvement of prediction accuracy to reach two-third of correct classification at 72h (63±3%). Interestingly, this dynamic collapsed after a memorisation phase of 24h with no longer exposure to sender plant: classification rates of LDA after 96h get back to random assignation (Fig. 5b).

**Figure 5:**
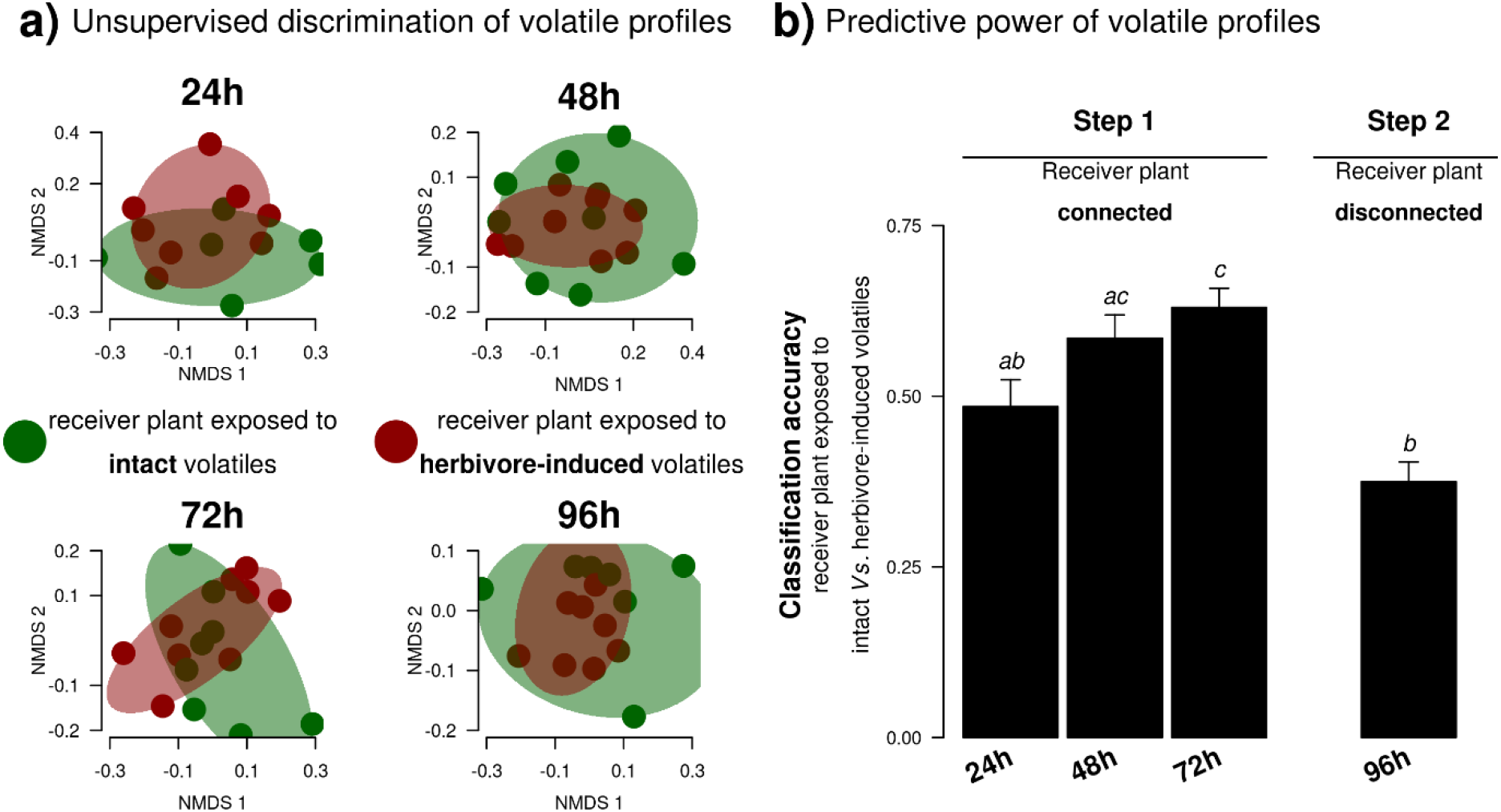
Volatile profiles emitted by receiver plants. **a)** Non-metric multidimensional scaling of volatile profiles was performed with Euclidean dissimilarity index. Stress values range from 0.112 (48h) to 0.152 (72h) **b)** Linear discriminant analysis of volatile profiles was used to predict whether receiver plants were exposed to intact or herbivore-induced sender plants (bootstrap resampling; n=100). Pairs of sender and receiver plants were connected during 72h. At 96h, receiver plants experienced a memory phase of 24h with no longer exposure to sender plants. Different letters indicate significant differences in correct classification rates of test datasets (pairwise Wilcoxon rank sum test; adjusted p.value <0.05)

### 3.3 Does chemodiversity help to understand information transfer?

Chemodiversity of VOC released by sender plants did not explain metabolomic chemodiversity collected on receiver plants (Fig 6a; linear models; *p.value*=0.843 and *p.value*=0.436 for plant pairs with intact and infested senders, respectively). However, we observed significant relationships between chemodiversity of VOC released by sender and receiver plants (Fig. 6b). The regression coefficient of the mixed model built with infested sender plants was higher (α=1.261; p.value<0.001) as compared to the model built with intact sender plants (α=0.472; p.value=0.001). A general mixed model pooling pairs of plants with intact and infested sender plants also indicate a significant relationship (p.value<0.001) and, more importantly, a significant interaction with the status of sender plant (p.value=0.004).

**Figure 6:**
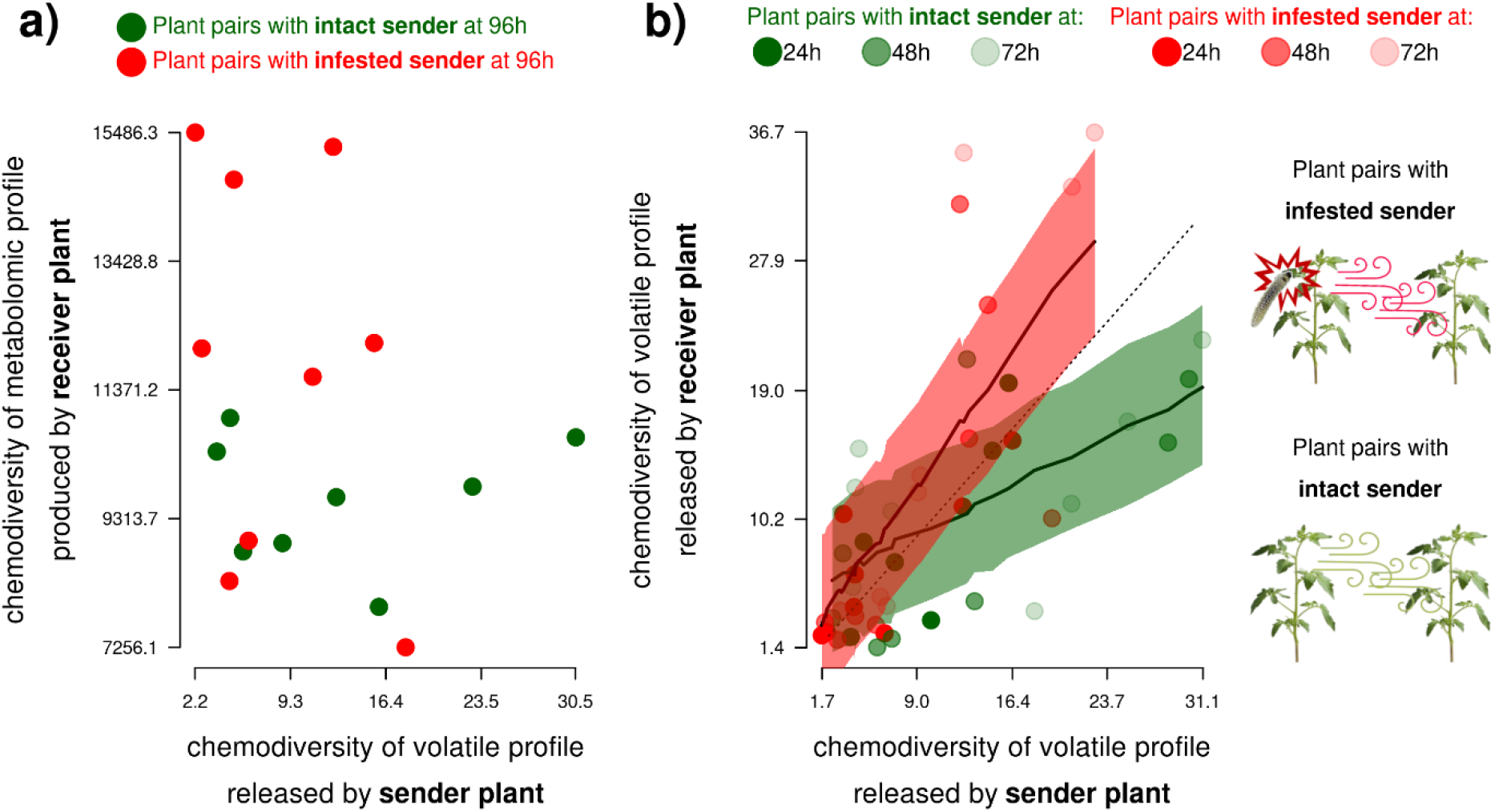
Chemodiversity among pair of sender and receiver plants. **a)** Relationships between chemodiversity of volatile profiles released by sender plant and chemodiversity of foliar metabolism of receiver plants collected at the end of the experiment (96h). **b)** Relationships between chemodiversity of volatile profiles released by sender and receiver plants all along plant-plant communication (24h, 48h and 72h). Chemodiversity is given as abundance-weighted functional Hill diversity index (diversity order q=1) computed from compound dissimilarity matrices (pairwise Jaccard distances between PubChem fingerprints of individual compounds). Solid lines indicate significant mixed linear models (pair of plants as random effect; Wald χ²; p.value ≤ 0.001). Transparency areas display prediction intervals of fixed effect (n=1000 simulations).

## 4. Discussion

This study takes benefit of tomato, i.e. a highly chemically-diversified agronomic model, to fill gaps in knowledge about mechanisms underpinning plant-plant communication. We highlight the interest of chemical ecology to integrate a chain of endogenous and exogenous processes leading to information transfer. Inside plants, we discuss the reprogramming of plant immune systems in both attacked sender plants and receiver plants exposed to HIPVs. In line with the concept of honest signal, our results point mirror effects with receiver plants aligning their defences on that of sender plants. Outside plants, in general, we observed positive correlations between chemodiversity of volatile profiles released by sender and receiver plants. However, herbivory in receiver plants significantly strengthen this relationship. Parallel to the mere composition of airborne cues, chemodiversity of volatile profiles opens new frontiers to understand information transfer among plants.

### 4.1 Plant immune system offers a mechanistic framework to explain the production of informative cues

Inside sender plants, metabolomic reprogramming induced by herbivores affects a large array of chemical classes involved in plant immune systems. Here, we captured changes in chemical classes underpinning herbivore recognition, signal transduction and defensive response (Howe and Jander, 2008; Snoeck et al., 2022). First, the general induction of amino acids and peptides may be related to the role of proteome in the recognition of herbivore-associated molecular patterns (aka HAMP, Vakhrusheva and Nedospasov, 2011). Second, changes of three fatty acid including a derivates of jasmonic acid (FT196) and phosphocholine (FT332) is probably linked with signal cascades connecting insect elicitors to defence response (for the role of lipid metabolism see Seth et al., 2024; for jasmonic acid Zhang et al., 2020; for phosphocholine Chen et al., 2018). Third, we can reasonably view the over-expression of terpenoids in general, and one alkaloid in particular, as part of the plant defences themselves since these metabolites play a key role in anti-herbivore resistance (Wari et al., 2022). In the specific case of terpenoids, we observed a re-allocation between two triterpenoid saponins with the down-regulation of FT489 and the up-regulation of FT530. Besides the defensive functions of terpenoid-related saponins (Osbourn, 1996), these compounds may also serve as intermediates of glycoalkaloids in *Solanaceaa* (Kohara et al., 2005). Further direct plant defences, the general up-regulation of terpenoids can also be considered as an important step in the production of airborne cues supporting plant-plant communication.

Indeed, monoterpene and sesquiterpenes offer good candidates to seek for info-chemicals conveying messages between plants since they support a high structural diversity (Rosenkranz et al., 2021). In tomato, terpenes are manufactured inside type VI trichomes with efficient storage capacities (Tissier et al., 2017). Soon after herbivore infestation, the rapid increase in monoterpene emissions could be viewed as a burst signal drawing on these reserves. Later, plants likely synthesize new pools of sesquiterpenoids as herbivory runs over time. Based on VOC comparisons after one-week of infestation, Anastasaki et al. (2018) demonstrated the induction of sesquiterpenoids, like β-caryophyllene or TMTT. Here, the kinetic of VOC profiles suggests that the balance between early-burst signals based on monoterpenes and delayed emissions of sesquiterpenes occurs relatively early, *i.e.* during the first 72h of infestation. This temporal pattern is cohesive with molecular works interested in the genetic machinery regulating *de novo* production of terpenoids (Tzin et al., 2015). In comparison to burst signals, Grandi et al. (2024) discuss how *de novo* signals arising from herbivore-specific VOC emissions could be of far most reliable interest for defence elicitation of receiver plants.

### 4.2 The response of receiver plants suggests that informative cues are reliable to stress

On the side of receiver plants, our results confirm plant-plant communication according to metabolomic reprogramming of tomato exposed to HIPVs. In this study, we tested the response of receiver plants after a ‘memorisation’ phase of 24h with no longer exposition to HIPV. Accordingly, the metabolomic response of receiver plants fits with priming rather than induction processes (Martinez-medina et al., 2016), even if further work accounting for pest performance is needed to conclude about potential links between plant-plant communication and defence elicitation in receiver plants. Nonetheless, our results show that direct herbivory and exposure to HIPV for sender and receiver plants, respectively, resulted in analogous metabolomic changes. More particularly, our result indicates that the responses of both sender and receiver plants rely on identical compounds among each of the three immunity layers. The peptide FT555, the conjugate of jasmonic acid FT196 and the terpenoids FT276 and FT530 were not only over-expressed in infested sender plants but also in intact receiver plant exposed to HIPV. Previous research pointed a convergence of chemical defences between sender and receiver plants (Grandi et al., 2024). In other words, plant-plant communication seems to operate as mirror effects with receiver plants matching up their physiology to that of sender plants. From an adaptive standpoint, this supports the idea that information conveyed between plants is explicit enough to warn receiver plant against appropriate stress.

### 4.3 Volatiles profiles released by receiver plants embed information larger than their mere composition

In contrast to metabolomic reprograming of receiver plant, the unsupervised analysis indicates that HIPV exposure does not affect the multivariate composition of VOC profiles released by receiver plants. In this line, (Catola et al., 2018) have shown that plant-plant communication in tomato result in minor changes of volatiles emitted by receiver plants. On the other hand, the supervised analysis points a cohesive pattern between the timeline of our two-step experimental setup and the latent ecological information embedded in VOC of receiver plants. During three days, as long as pairs of plants remained connected, we observed a progressive increase in predictive power of VOC profiles to classify receiver plants exposed to intact volatile or herbivore-infested plants. But, as soon as pairs of plants were disconnected, this predictive power collapsed. This responsiveness within the dynamic of predictive power of VOC released by receiver plant is likely intrinsic to such plants since we systematically purged airflow from sender plants. Hence, supervised analysis of volatile profiles yields a particular facet of response of receiver plant that provide information about exposition to airborne cues. Beyond the mere composition of individual VOC, and among the latent patterns captured by supervised analyses, we can reasonably think that group-level properties of volatile profiles released by receiver plant such as compound richness, evenness or structural disparity may contribute to their predictive power.

### 4.4 Chemodiversity of volatile profiles as a new frontier of information transfer

Based on these findings, we investigated the chemodiversity of volatile profiles released by specific pairs of sender and receiver plants. Here again, we observed a mirror effect. The more chemodiversity of VOC profiles of sender plants increases, the more VOC profiles of receiver plants are diversified. However, herbivory on sender plants strengthen this linear relationship. This suggests that chemodiversity itself could fulfil a role in the propagation of information between plants. In opposition to animal systems, redundancy and semi-autonomous modularity of plant sensory systems are generally considered as less sensitive to directed cues (Karban, 2021). This architecture at the plant level could be appropriate to capture chemodiversity of complex volatile cues. At the molecular level, current understanding of VOC uptake is of valuable interest to discuss the respective roles of composition and chemodiversity of volatile profiles in plant-plant communication.

Recent reviews categorize plant volatile uptakes along a gradient from active to passive processes (Arimura and Uemura, 2025; Kessler et al., 2023; Wang and Erb, 2022). Active routes of VOC uptakes support the idea that the response of receiver plants depends mainly on the detection of specific compounds via ligand-receptor interactions (see Meents and Mithöfer, 2020 for analogy of HIPV with damage-associated molecular patterns). Thereby, the presence of particular HIPV, rather than chemodiversity, could be responsible for the activation of entire defensive pathways. This may explain why chemodiversity of VOC profiles released by sender plant is not related to chemodiversity of metabolomic make-ups inside receiver plants. Among HIPV, different clues converge to point sesquiterpenoids like β-caryophyllene as good candidates. First, while the perception of this compound is known to orchestrate plant signalling networks (Frank et al., 2021), β-caryophyllene has also played a key role in the discovery of the first VOC-binding proteins regulating gene expression on plant model belonging to *Solanaceae* (Nagashima et al., 2019). Second, while constitutive emissions of sesquiterpenoids remains scarce in intact tomato (Amegan et al., 2024), our results show that β-caryophyllene displayed the highest increase in response to herbivory. A low basal level of emission along with high inducibility make it a good candidate as infochemical.

In contrast, passive VOC uptakes suggest that volatile infochemicals can be adsorbed in waxy cuticles, enter plants and pass cell wall before to trigger a response via access to cytoplasm or membrane depolarization. It is reasonable to expect that such passive mechanisms are less susceptible to elicit targeted pathway based on orchestrated response since they do not rely on particular signal recognition occurring through ligand-receptor interaction. Instead, passive VOC uptakes are expected to affect concomitantly different pathway but in a more versatile manner as compared to active mechanisms. In our case, this may explain why the links between chemodiversity of VOC profiles released by sender and receiver plants does not persist after the pairs of plants have been disconnected (see 4.3). So far, while plant-plant communication is generally thought at relatively small distances, c.a 1m (Heil and Adame-Álvarez, 2010), the role of chemodiversity as carrier of information could open new avenues to study “waves” of communications in patch of intact plants distributed along gradients from herbivore-infested plants.

## 5. Conclusion and perspectives

To sum up, we have shown that tomato provides a good model to address a set of mechanisms underpinning plant-plant communication. This study brings cohesive understanding about production, emission and cues perception. On the side of sender plant, herbivore attacks induce the entire immune system and cause important shifts in volatile emissions to the benefit of sesquiterpenoids, like β-caryophyllene, that potentially convey information. On the side of receiver plant, metabolism reprogramming confirm cues detection although entomological bioassays are now required to ensure that this response is part of defence priming.

Analogously to mirror effects, direct herbivory and exposure to HIPV on sender and receiver plants, respectively, result in similar patterns regarding both the plant immune system and the chemodiversity of volatile emissions. Hence, our results suggest that informative cues driving plant-plant communications are explicit enough to warn neighbouring plants against appropriate stress. So far, the concept of ‘honest’ signal in communication systems has been mainly studied in plant-pollinator interactions (Knauer et al., 2021). We need a surge in interest to understand the determinants of honesty in plant-plant communications (e.g., nature of stress experienced by sender plant, physiological status of receiver plant, integration of multiple cues).

Moreover, this study paves interesting perspectives to reconsider informative channels underpinning plant-plant communications. Besides individual info-chemicals, our results suggest that chemodiversity of volatile profiles may contribute to information transfer. Further work based on experimental manipulation of volatile profiles is needed to investigate joint effects of the composition and the chemodiversity of airborne cues. At this point, it remains unclear how sensing system of plants could take into account defined ranges of volatile chemodiversity with threshold values informing about when it becomes relevant for plant-plant communication. In the near future, if the role of chemodiversity as vehicle of information is confirmed, it could open promising avenues to better understand the efficiency of communication systems in noisy (Wilson et al., 2015) or perturbated environments (Sun et al., 2026).

## Supporting information

TabS1

Dataset of volatile emission

Dataset of leaf metabolomic

Metadata related to volatile collection

Metadata related to leaf sampling

R script used for deconvolution of volatile emissions

R script used for deconvolution and dereplication of leaf metabolomic

R script used for statistics

## 6. Acknowledgements

This work was supported by the Metaboscope, part of the “Platform 3A” funded by the European Regional Development Fund, the French Ministry of Research, Higher Education and Innovation, the region Sud Provence-Alpes-Côte d’Azur, the Departmental Council of Vaucluse and the Urban Community of Avignon. We are grateful to Vanessa Kuentz for her involvement in the coordination, the maintenance and the functioning of the metabolomic platform.

## Supplementary Material

**Figure S1:**
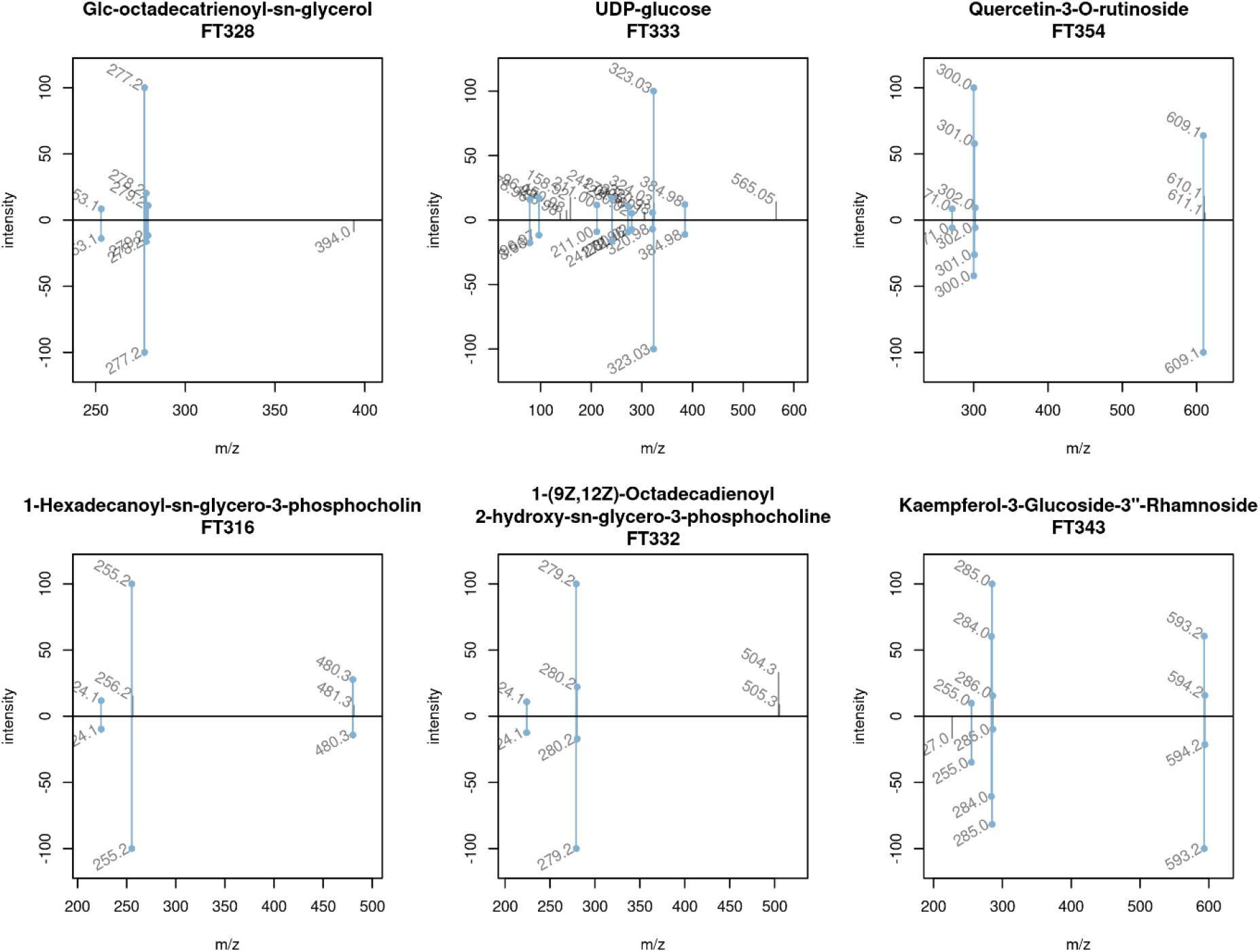
Annotation of foliar metabolomic features based on MS2 spectra. Pairs of experimental (positive intensities) and reference spectra (negative intensities) retrivied from MassBank (release 2024.11)

**Figure S2.**
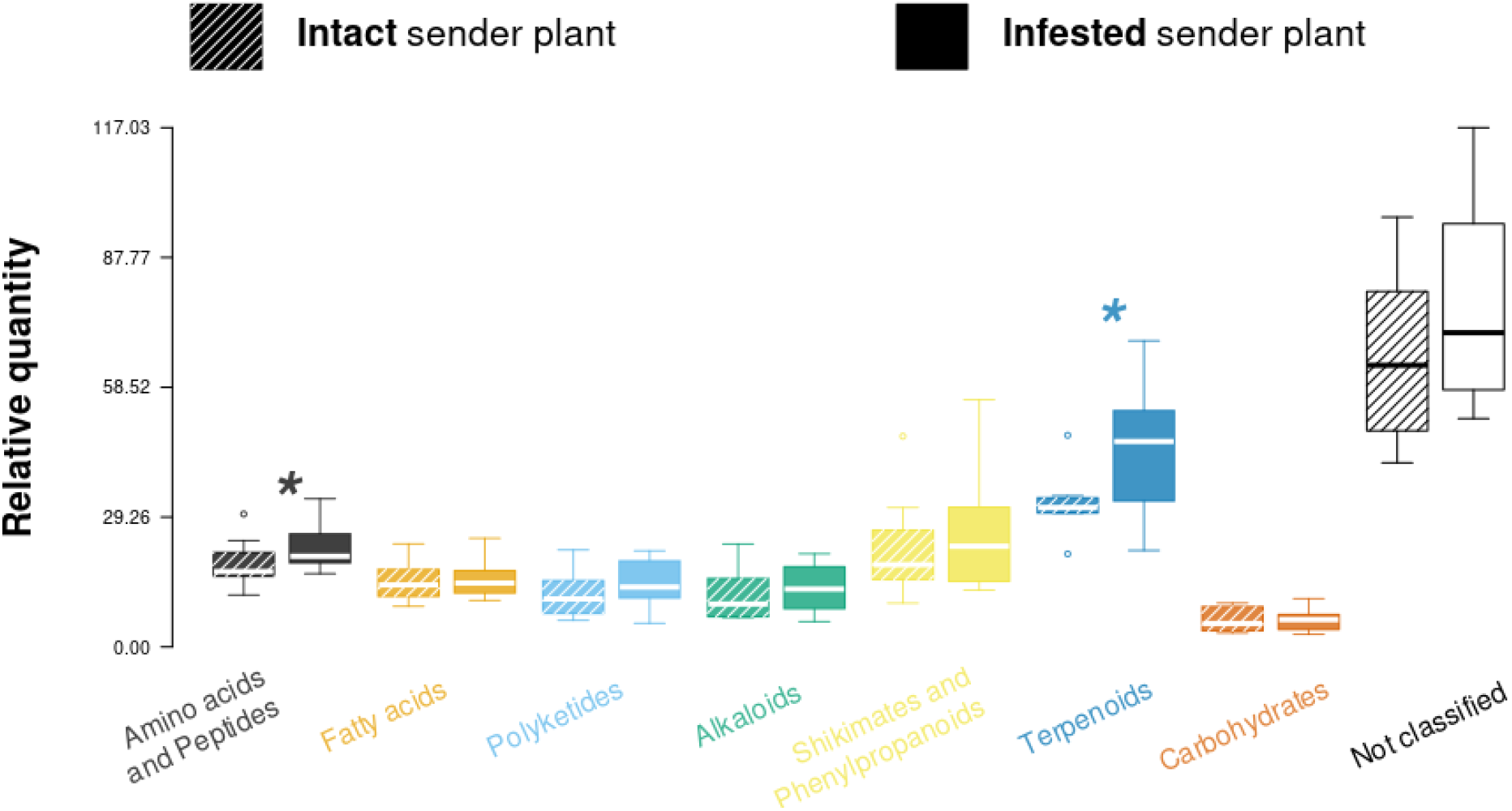
Pathway enrichment of sender plants. Asterisk indicates significant differences in total quantity of metabolomic features assigned to each pathway (one-tailed Wilcoxon rank sum test; * : *p.value* <0.05)

**Figure S3.**
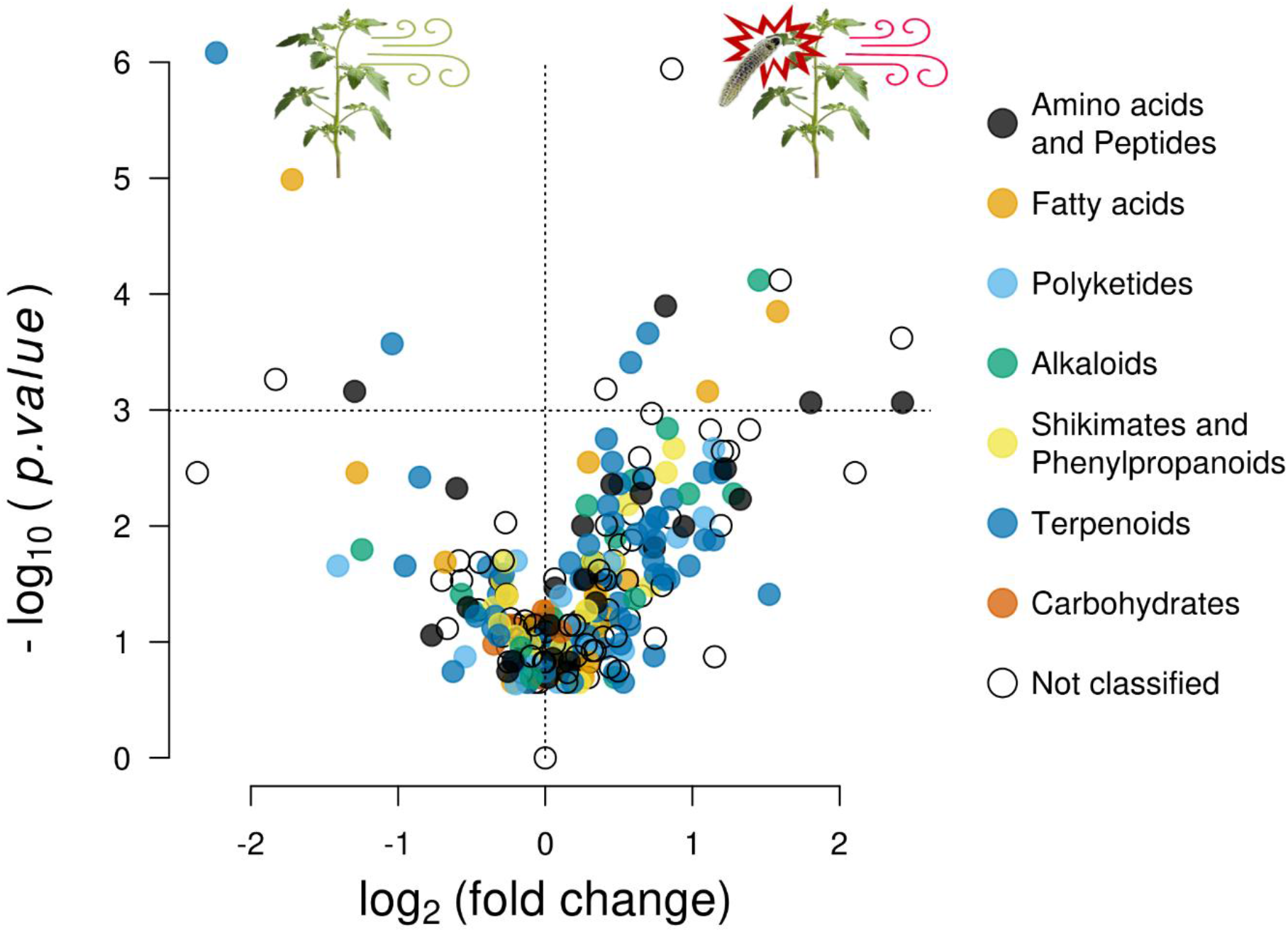
Volcano plot of sender plants. The 291 metabolomic features detected are coloured according to chemical pathway (NPCclassifier ontology). The vertical line separates feature over-expressed in infested sender plant (positive values on x-axes) as compared to intact sender plant (negative values on x-axes). Features with significant differences are displayed above the horizontal line (one-tailed Wilcoxon rank sum test; *p.value* < 0.05).

